# RNA polymerase III transcription-associated polyadenylation promotes the accumulation of noncoding retrotransposons during infection

**DOI:** 10.1101/2025.05.14.654096

**Authors:** Azra Lari, Sahil B. Shah, Xiaowen Mao, Priyanka Sanghrajka, John Karijolich, Liana F. Lareau, Britt A. Glaunsinger

**Author notes:** A.L and S.B.S contributed equally to this work.

## Abstract

The accumulation of RNA Polymerase III (Pol III) transcribed short interspersed nuclear element (SINE) retrotransposon RNA is a hallmark of various cellular stressors, including DNA virus infection. However, the molecular mechanisms driving the induction of these normally repressed loci are largely undefined. Here, we reveal that in addition to Pol III transcriptional induction, gammaherpesvirus infection stimulates mRNA-like 3’ end processing of SINE RNAs that leads to their stabilization. We developed a convolutional neural network (CNN)-based model that identified a polyadenylation-associated motif as the key hallmark of infection-induced SINEs. Indeed, mRNA polyadenylation machinery is recruited in a Pol III-dependent manner to virus-induced loci, including B2 SINE and tRNA genes. Infection causes enhanced polyadenylation of SINE ncRNA, which is required for its stable accumulation. This virus-host interaction therefore highlights an inducible, coupled relationship between Pol III transcription and mRNA-like polyadenylation. It also reveals that co-option of the polyadenylation machinery by Pol III is a mechanism to increase the abundance of noncoding RNA during pathogenic stress.

**SIGNIFICANCE:** Short interspersed nuclear elements (SINEs) are hyperabundant and transcribed by RNA polymerase III (Pol III) to produce noncoding retrotransposons. Although generally not detectable in healthy somatic cells, SINE RNA expression is upregulated during stress, including viral infection and inflammatory diseases. We used gammaherpesvirus infection to uncover pathways leading to increased SINE RNA expression. Using a newly developed deep learning model and genomics analyses, we reveal that infection-induced accumulation of SINEs is driven by increased Pol III transcription and Pol III-dependent recruitment of polyadenylation machinery. This stimulates polyadenylation of SINEs, which is a known stabilizer of these noncoding transcripts. Our findings suggest that inducible alterations to Pol III transcript 3’ end processing modulate the abundance of noncoding retrotransposons during pathogenic stress.

## INTRODUCTION

Transposable elements have been found in nearly all sequenced eukaryotic genomes (1). In mammals, retrotransposons constitute the largest group among these parasitic DNA elements and are capable of propagating and integrating throughout the genome via RNA intermediates (2). Approximately 10% of the mammalian genome consists of short interspersed nuclear elements (SINEs), a class of retrotransposons whose evolutionary origins trace back to endogenous RNA polymerase III (Pol III) transcripts. Human Alu elements and the murine B1 SINEs are derived from the 7SL RNA, a component of the signal recognition particle, while the murine B2 SINE family is derived from tRNA genes (3–6).

Even when not independently transcribed, SINEs can impart gene regulatory functions, including when they are ‘embedded’ within protein-coding genes or have been exapted as enhancer or promoter elements (7–13). However, when transcribed by Pol III, they produce short (<500 bp), noncoding RNAs (ncRNAs). SINE retrotransposition requires the co-option of the reverse transcriptase machinery encoded by long interspersed nuclear elements (LINEs), which recognize a polyadenosine (polyA) tail on the SINE transcript (14–16). Some SINEs, like Alu elements, contain templated poly(A) sequences. Others, including a subset of murine B2 SINEs, instead contain a poly(A) signal sequence (PAS) upstream of the Pol III terminator sequence and recruit the messenger RNA (mRNA) polyadenylation machinery at an unknown stage of their biogenesis (5, 17–21). SINE retrotransposition can cause insertional mutagenesis (22), and thus, most of them are repressed in healthy somatic cells, including through trimethylation of lysine 9 on histone H3 (H3K9) marks and CpG methylation (23–25).

Even in the absence of retrotransposition, accumulation of Pol III transcribed SINE ncRNAs has gene regulatory and pathologic potential. In particular, the accumulation of SINE ncRNA is a hallmark of various cellular stressors and inflammatory-associated diseases (26–34); indeed, Alu ncRNAs were the first endogenous nucleic acid shown to activate the NLRP3 inflammasome (29). B2 SINE ncRNAs are rapidly upregulated in response to thermal stress, where they can bind RNA polymerase II (Pol II) and repress preinitiation complex formation, as well as suppress transcription elongation of heat shock-inducible genes (35–39). Viral infection is also a prominent inducer of SINE ncRNAs, which can influence innate immune signaling and alter cellular and viral gene regulation. Infection with several DNA viruses, including adenovirus, herpes simplex virus (HSV-1), and the model murine gammaherpesvirus MHV68, lead to a dramatic accumulation of SINE ncRNAs (40–46). Recent genome-wide studies in herpesvirus-infected cells demonstrated widespread increases in the expression of thousands of murine B2 SINE loci as well as tRNA genes, both of which use internal type II Pol III promoters (47–49). Pol III induction during viral infection is largely specific to loci with type II promoters, as Pol III loci with type I (e.g., 5S rRNA) or III (e.g., U6, 7SK) promoters are minimally impacted. Several individual viral proteins have the potential to increase SINE expression, partly through their mitogenic signaling activity (50, 51). However, the mechanisms driving the pronounced accumulation of SINE ncRNAs during infection remain largely unknown.

Here, we sought to define the mechanisms driving B2 SINE ncRNA accumulation during viral infection, using the herpesvirus MHV68 as a model. We found that increased Pol III occupancy only partially explained the B2 SINE induction during infection. We therefore developed a convolutional neural network (CNN) based model, which revealed that an mRNA-like polyadenylation-associated motif is the primary discriminating sequence feature of infection-induced B2 SINE genes. We confirmed the requirement for mRNA polyadenylation machinery for both B2 SINE ncRNA polyadenylation and accumulation in infected cells. Furthermore, we discovered that infection stimulates Pol III-dependent recruitment of mRNA CPSF components to sites of B2 SINE transcription, suggesting inducible, co-transcriptional polyadenylation of B2 SINE ncRNAs. Notably, CPSF recruitment to Pol III type II promoters is not restricted to murine genomes but also occurs at human SINE and tRNA genes. Thus, the noncanonical use of mRNA polyadenylation machinery to influence ncRNA transcript abundance may be a conserved feature of Pol III transcription.

## RESULTS

### RNA Polymerase III exhibits enhanced binding to a subset of type II promoters during MHV68 infection

Approximately 30,000 Pol III transcripts from loci with type II promoters increase in abundance upon MHV68 infection of mouse fibroblasts (47, 48). To test the extent to which Pol III transcriptional induction underlies the increased levels of these RNAs, we performed chromatin immunoprecipitation experiments followed by sequencing (ChIP-seq) to map the genome-wide occupancy of the major catalytic subunit of Pol III, Polr3a, in mock- and MHV68-infected mouse NIH3T3 fibroblasts. Given the repetitive, degenerate, and arrayed nature of most Pol III transcribed genes, we applied stringent mapping criteria and called peaks using MACS2 (52). We detected 7,241 additional Polr3A-occupied genes in MHV68-infected cells than in mock-infected cells (**Fig. 1A, Fig. S1A**). As expected, we also detected Polr3A binding to the Pol III promoter-containing genes on the MHV68 genome (termed tRNA-miRNA-encoding RNAs (TMERs) (53)) (**Fig. S1B**).

**Figure 1.**
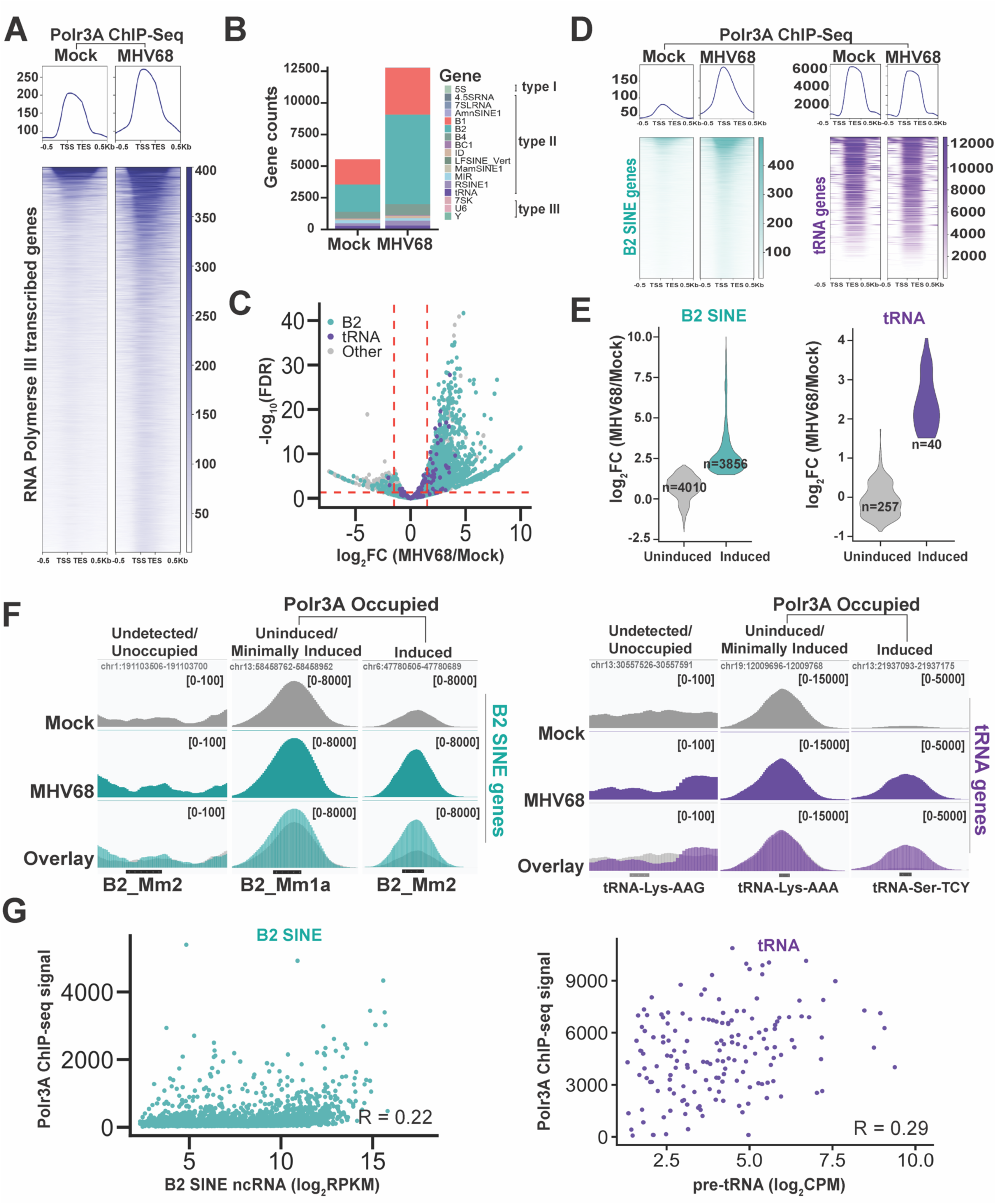
RNA Polymerase III binding to the cellular genome during MHV68 infection (A) The top panel is a metagene plot displaying Polr3A ChIP-seq signal across all Polr3A peaks associated with Pol III transcribed genes in mock-treated or MHV68-infected NIH3T3 cells (n=14739). Alignment files were averaged from two replicates. ChIP-seq signal was plotted as a histogram with 10 bp bins for −500 to +500bp around the transcription start site (TSS) with 20 bins/gene. In the bottom panel, each row of the heat map corresponds to the ChIP-seq signal from each Pol III gene from the metaplot above. Heat maps are ranked by Polr3A ChIP-seq coverage values. (B) The stacked bar graph shows the number of genes within each category of Pol III transcribed genes annotated by RepeatMasker (93) that Polr3A peaks identified overlap within mock-treated or MHV68-infected samples. The promoter types (I, II, and III) associated with Pol III genes is shown. (C) DiffBind (54) was used to perform differential binding analysis of Polr3A occupancy data at regions overlapping with Pol III genes from mock-treated and MHV68-infected samples. The values plotted are log_2_ fold change (FC) (MHV68 vs. Mock) versus the false-discovery rate (FDR) *p*-value. A *p-value* threshold of 0.05 was used. Dashed red lines denote significant p-values and ≤-1.5 log2FC ≥1.5. (D) The top panels are metagene plots displaying ChIP-seq signal across all B2 SINE genes (left, n=7472) or tRNA genes (right, n=299) occupied by Polr3A. Alignment files were averaged from two replicates. ChIP-seq signal was plotted as a histogram with 10 bp bins for −500 to +500bp around the TSS with 30 bins/gene. In the bottom panel, each row of the heat map corresponds to the ChIP-seq signal for each B2 SINE or tRNA gene from the metaplot above. Heat maps are ranked by Polr3A ChIP-seq coverage values. (E) Log_2_FC values from the differential binding analysis in panel (C) for B2 SINE (left) and tRNA (right) genes. *Uninduced* is defined as an FDR>0.05, or 0<log_2_FC<1.5 (FDR≤0.05), or 0>log_2_FC>-1.5 (FDR≤0.05) and *induced* is defined by log_2_FC≥1.5 (FDR<0.05). The plot displays the number of genes (n) associated with each category. (F) Polr3A ChIP-seq coverage across select B2 SINE and tRNA genes from mock-treated or MHV68-infected samples. Alignment files were averaged from two replicates and visualized in the Integrative Genome Viewer (IGV) (96). Y-axis maximum and minimum values are within brackets. Loci shown were selected based on the differential binding analysis from panel (C), where *unchanged or minimally induced* is defined as an FDR>0.05, or 0<log_2_FC<1.5 (FDR≤0.05), or 0>log_2_FC>-1.5 (FDR≤0.05) and *induced* is defined by log_2_FC≥1.5 and FDR<0.05. (G) ChIP-seq signals associated with Polr3A occupied peaks at B2 SINE (left) and tRNA (right) genes from panel (A) during infection were plotted against RNA expression data from previously published datasets in MHV68-infected mouse fibroblasts(47, 48). The correlation (R) between ChIP-seq signals and RNA-expression data is denoted on the graphs and was calculated using the Pearson correlation coefficient.

On the cellular genome, Polr3A peaks overlapped with all three Pol III promoter types in both mock and infected cells (**Fig. 1B**). Polr3A differential binding analysis using DiffBind (54) confirmed that infection-associated increases in Polr3A occupancy at Pol III genes primarily occurred on type II promoters, this included B2 SINE and tRNA genes (**Fig. 1 C and D and Dataset S1**). Notably, B2 SINE genes made up the greatest number of differentially bound loci when compared to all other Pol III-transcribed genes, whereas Polr3A binding to a majority of tRNA genes remained largely unaltered between conditions (**Fig. 1C**). Overall, genes with type I (e.g., 5S rRNA) or type III (e.g., U6 RNA, 7SK RNA) promoters showed minimal or reduced Polr3A binding when comparing MHV68-infection to mock conditions, consistent with previous work (**Fig. S2 A and B**) (45, 47, 48, 50, 51).

We conclude from the above data that there is likely increased Pol III transcription at thousands of type II promoters upon MHV68 infection. However, several observations suggest that additional mechanisms beyond increased Pol III binding contribute to the abundance of these RNAs. Nearly half of the Polr3A occupied B2 SINE genes, and the majority of occupied tRNA genes had modest to no induction of Polr3A occupancy upon MHV68 infection (**Fig. 1 E and F**). Additionally, our prior B2 SINE-specific sequencing (SINE-seq) data (47) identified infection-induced increases in B2 SINE ncRNAs from more than twice the number of loci than those with increased Polr3A occupancy detected here. We found only a weak correlation between the Polr3A ChIP-seq signals and previously published B2 SINE ncRNA or pre-tRNA expression levels in MHV68-infected mouse fibroblasts (47, 48) **(Fig. 1G).** We therefore hypothesized that there may be sequence features specific to the induced loci that promote the accumulation of their RNA in infected cells via a mechanism other than increased Pol III binding. We first searched for such features on B2 SINE loci, as these comprise the vast majority of the infection-induced Pol III transcripts.

### Infection-responsive B2 SINE loci contain sequences that promote polyadenylation

Prior SINE-seq data in NIH3T3 cells showed that the highest accumulating B2 SINE transcripts fall within the SINE families B2_Mm1a, B2_Mm1t, and B2_Mm2 (47). We therefore designed a a convolutional neural network (CNN) to identify sequences within the B2 SINE gene body in these families that distinguish infection-induced from uninduced B2 SINE loci. This binary classification CNN, termed <u>S</u>equence <u>A</u>nalysis of <u>M</u>urine <u>B</u>2 <u>A</u>ccumulated <u>R</u>NAs-Net (SAMBAR-Net), enabled base pair-level resolution of sequences linked to B2 SINE ncRNA expression during MHV68 infection. SAMBAR-Net is based on the DenseNet CNN architecture (55), which provides the advantages of parameter efficiency and compactness. We reasoned that B2 SINE ncRNAs would only accumulate if they contained intact internal promoter sequences and other regulatory motifs, provided the locus was in a chromatin-accessible region. Thus, the input data used for CNN training focused on internal B2 SINE gene sequences and was labeled based on each B2 SINE gene’s expression level, which was obtained from our previously generated SINE-seq dataset (47). We focused on the families containing the greatest number of expressed B2 SINE genes and excluded the B3 and B3A B2 SINE families since they were severely underrepresented. Across all three B2 SINE families analyzed, SAMBAR-Net effectively predicted whether a B2 SINE locus would be expressed, with area under the ROC curve (AUC) values of 0.85, 0.91, and 0.97 for the B2_Mm1a, B2_Mm1t, and B2_Mm2 families, respectively (**Fig. 2A**). When analyzing our model’s gradients using Transcription Factor Motif Discovery from Importance Scores (TF-MoDISco) (56), an algorithm used for identifying motifs from base pair-level importance scores, we were able to identify specific TGT-containing subsequences within a known τ motif, which encodes a sequence that enhances polyadenylation through binding of the mammalian cleavage factor I complex (CFIm) (57, 58) (**Fig. 2B**). This prediction is in line with studies showing that the τ motif encoding a UGUA RNA sequence, preceding a canonical mRNA-like hexameric (AAUAAA) polyadenylation signal sequence (PAS), is necessary for CFIm25 (murine Nudt21) binding to the newly synthesized RNA and forming the full polyadenylation complex on both mRNA and B2 SINE ncRNAs (17–20, 57, 58). Thus, the most important discriminating factors for highly expressed B2 SINE genes in MHV68 infected cells are sequences that promote polyadenylation.

**Figure 2.**
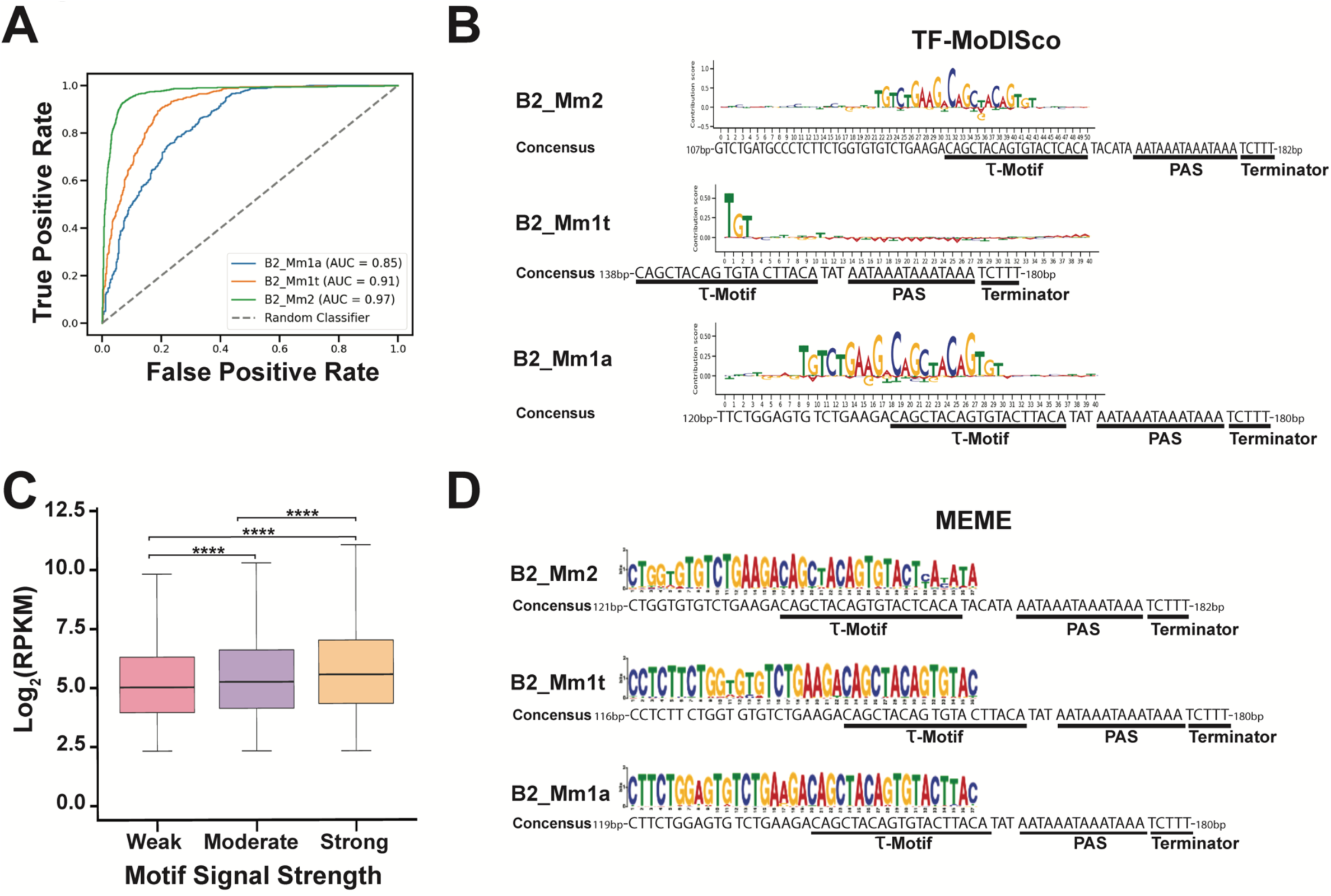
Infection-responsive B2 SINE loci contain sequences that promote polyadenylation (A) ROC curve of the SAMBAR-Net CNN across all three B2 SINE families analyzed. AUC values for each B2 SINE family measured how well SAMBAR-Net performed in discriminating between expressed (RPKM ≥10) and unexpressed (RPKM = 0) B2 SINE DNA sequences from previously published SINE expression datasets (47). (B) TF-MoDISco analysis was performed on SAMBAR-Net to identify the most prominent predictive sequence features of expressed versus unexpressed B2 SINE families. The LOGO plots shown display the top-ranked motifs for each B2 SINE family analyzed. B2 SINE consensus sequences (25) for each family are matched to the nucleotide position for each LOGO plot, highlighting the τ motif, polyadenylation signal sequence (PAS), and Pol III terminator sequence. (C) Boxplot of log_2_RPKM values for expressed B2 SINE genes (RPKM ≥5) during MHV68 infection (47) partitioned into three quartiles by PWMScores. Weak, moderate, and strong labels correspond to increasing PWMScores, which measure the prediction strength of a motif. Error bars show the standard deviation (SD), and statistics were calculated using the Mann-Whitney U test. ****, p-value <10e-17. (D) Discriminative MEME (60) analysis of expressed (RPKM ≥10) versus unexpressed (RPKM = 0) B2 SINE DNA sequences identified from previously published SINE-expression dataset in MHV68 infected NIH3T3 cells (47). LOGO plot shows the top-ranked discriminatory DNA sequence motif for each B2 SINE family analyzed. B2 SINE consensus sequences (25) for each family are matched to the nucleotide position for each LOGO plot, highlighting the τ motif, PAS, and Pol III terminator sequence.

To evaluate the relationship between B2 SINE ncRNA accumulation and the discriminatory sequence identified through TF-MoDISco, we generated a position weight matrix (PWM) to identify B2 SINE sequences containing the polyadenylation-associated motif. Since many B2 SINE genes have acquired mutations, we wanted to determine whether weaker predicted motifs (i.e., more divergent sequences) had a negative impact on B2 SINE ncRNA accumulation. Using PWMScan (59) to obtain PWMScores, a metric that relates to the prediction strength of a motif, for each expressed B2 SINE locus during MHV68 infection, we found that ∼99% of expressed B2 SINE genes contained the motif of interest (27572 of the 27943 total expressed B2 SINEs identified in (47)). We also observed a positive correlation between the strength of the prediction for the motif and B2 SINE ncRNA abundance (**Fig. 2C**). These data suggest that mutations within the motif, and thus weaker polyadenylation signals, negatively impact B2 SINE expression levels.

As an orthogonal method to validate SAMBAR-Net’s findings, we applied a discriminative DNA sequence motif discovery algorithm (MEME) (60). Although MEME lacks the base-pair level resolution of SAMBAR-Net, it also identified a TGT-rich sequence preceding the PAS as the top-ranked motif found in the majority of induced but not uninduced B2 SINE genes (**Fig. 2D**). Thus, although expressed B2 SINE loci have canonical Pol III termination sequences, both approaches implicated polyadenylation as a key feature of infection-induced accumulation of these retrotransposon RNAs. This hypothesis is further bolstered by data from uninfected cells demonstrating that a subset of SINEs can be polyadenylated by the mRNA polyadenylation machinery in a manner dependent on the PAS and τ motif (17, 19, 20, 61).

### B2 SINE RNA polyadenylation during MHV68 infection is dependent on mRNA cleavage and polyadenylation factors

We examined the polyadenylation status of B2 SINE ncRNAs in mock and MHV68-infected NIH3T3 cells by northern blotting total RNA with a B2 SINE probe (**Fig. 3A**). The increased levels of B2 SINE ncRNAs in MHV68-infected cells comprised a mixture of a short band, which is the expected length of the short non-polyadenylated species, and a longer smear. Deadenylation of the B2 SINE ncRNA by pre-treatment with oligo-deoxythymidine (Oligo dT) and RNase H collapsed the smear to the short band (**Fig. 3A**), confirming that a significant fraction of the virus-induced B2 SINE ncRNAs are polyadenylated.

**Figure 3.**
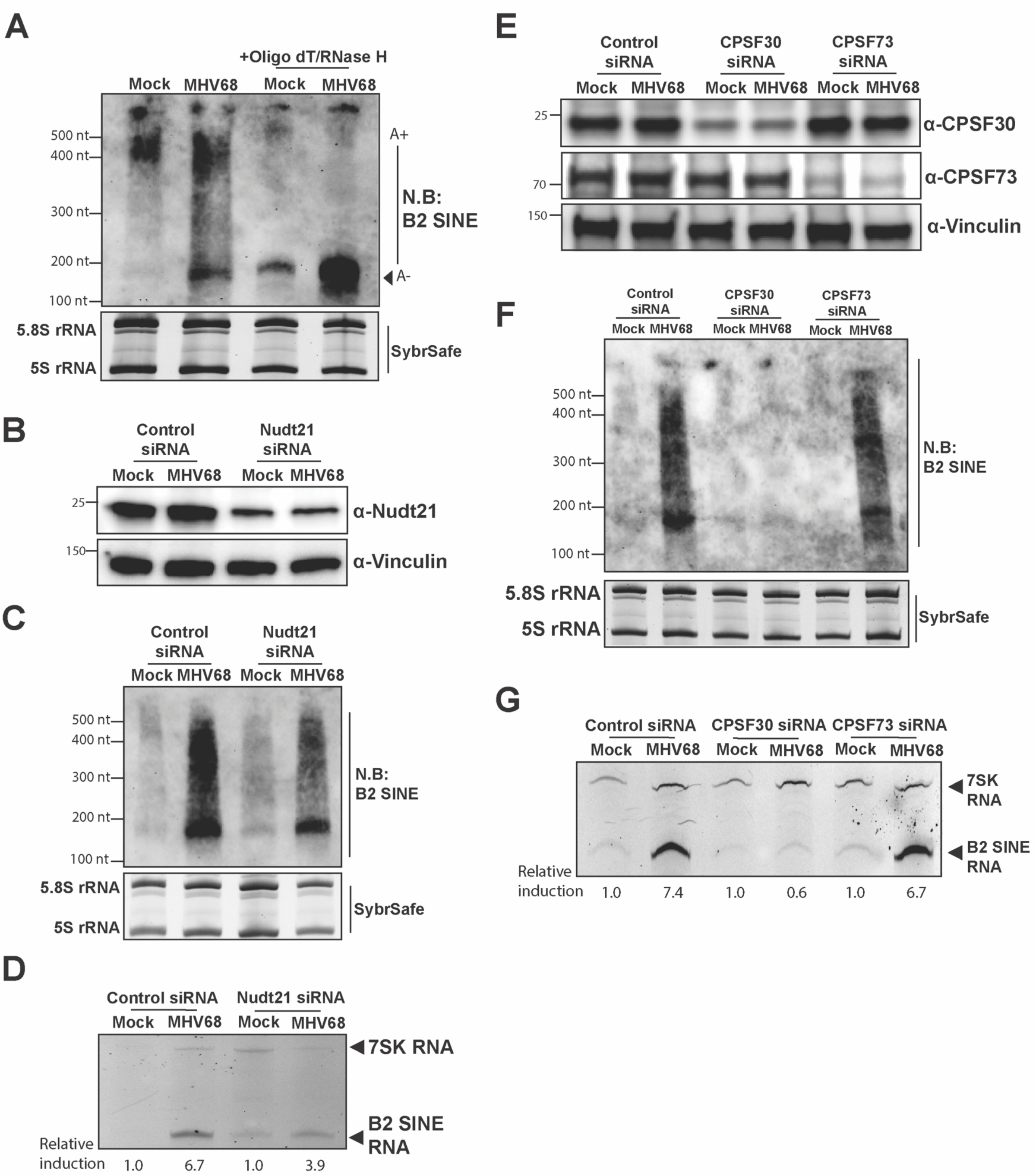
B2 SINE RNA polyadenylation during MHV68 infection requires mRNA cleavage and polyadenylation factors (A) NIH3T3 cells were mock-treated or infected with MHV68 at an MOI of 5. At 24 hpi, cells were harvested and lysed to extract total RNA. RNA was then digested with RNaseH in the presence or absence of oligo(dT) and subjected to Northern blotting using a primer specific to B2 SINE ncRNAs. SybrSafe stained gel shows 5.8S and 5S rRNA species in the bottom panel as loading controls. (B) NIH3T3 cells transfected with control non-targeting, or Nudt21 targeting siRNAs were mock-treated or infected with MHV68 at an MOI of 5. At 24hpi, cells were harvested and lysed to extract total protein and were analyzed by Western blotting with antibodies against Nud21and Vinculin (loading control). (C) Cells from panel B were also harvested for total RNA extraction and subjected to Northern blotting using a primer specific to B2 SINE ncRNAs. SybrSafe stained gel shows 5.8S and 5S rRNA species in the bottom panel as loading controls. (D) Total RNA extracted in panel C was also used for primer extension using primers specific to B2 SINE ncRNAs or 7SK RNA (control). The relative induction of B2 SINE ncRNAs for each sample was measured as the ratio of the mean integrated intensity between 7SK RNA and B2 SINE ncRNA levels and normalized to the mock-treated controls. (E) NIH3T3 cells transfected with control non-targeting, CPSF30 targeting, or CPSF73 targeting siRNAs were mock-treated or infected with MHV68 at an MOI of 5. At 24hpi, cells were harvested and lysed to extract total protein and were analyzed by Western blotting with antibodies against CPSF30, CPSF73, and Vinculin (loading control). (F) Cells from panel E were also harvested for total RNA extraction and subjected to Northern blotting using a primer specific to B2 SINE ncRNAs. SybrSafe stained gel shows 5.8S and 5S rRNA species in the bottom panel as loading controls. (G) Total RNA extracted in panel F was also used for primer extension using primers specific to B2 SINE ncRNAs or 7SK RNA (control). The relative induction of B2 SINE ncRNAs for each sample was measured as the ratio of the mean integrated intensity between 7SK RNA and B2 SINE ncRNA levels and normalized to the mock-treated controls.

Nudt21 (human CFIm25) recognizes a UGUA sequence encoded within the ! motif that promotes the assembly of the polyadenylation complex. Therefore, if the SAMBAR-Net prediction was correct, Nudt21 should be required for infection-induced B2 SINE ncRNA polyadenylation. Indeed, siRNA-mediated reduction of Nudt21 levels resulted in a decrease in B2 SINE ncRNA polyadenylation during MHV68 infection as compared to samples treated with control non-targeting siRNAs (**Fig. 3 B and C**). The reduced polyadenylation of B2 SINE ncRNA in Nudt21 knock-down conditions decreased their overall abundance, as quantified by primer extension analysis of total B2 SINE ncRNA (**Fig. 3D**).

As a component of the CFIm complex, Nudt21/CFIm25 aids in the recruitment of other polyadenylation factors like the CPSF machinery to pre-mRNAs (57, 58). We therefore next investigated the role of the mRNA CPSF complex in infection-induced B2 SINE ncRNA polyadenylation. Using siRNA-mediated knockdowns, we depleted CPSF30, a factor required for PAS recognition and cleavage site determination, and CPSF73, the RNA endonuclease that cleaves nascent RNA (**Fig. 3E)**. Depletion of CPSF30 prevented B2 SINE ncRNA polyadenylation during infection, but depletion of CPSF73 had minimal effect on polyadenylation relative to the control siRNA (**Fig. 3F**). Primer extension analysis of total B2 RNA levels also confirmed that depleting CPSF30 but not CPSF73 prevented infection-induced SINE ncRNA accumulation during infection (**Fig. 3G**), further suggesting that polyadenylation, which is known to stabilize B2 SINE ncRNA (21), promotes the increased abundance of these Pol III transcripts during MHV68 infection. The requirement for CFIm25 and CPSF30 but not CPSF73 is consistent with previous studies using plasmid-encoded SINEs in uninfected cells (20), suggesting the machinery involved is similar in uninfected and MHV68-infected cells. However, infection greatly stimulates B2 SINE ncRNA polyadenylation relative to unperturbed conditions.

### MHV68 infection drives recruitment of mRNA cleavage and polyadenylation factors to B2 SINE loci

Pre-mRNA processing and polyadenylation are coupled to Pol II transcription. The CPSF complex is thought to be initially recruited to the 5′-end of the gene, possibly by transcription factor complex TFIID, then handed off to the c-terminal domain (CTD) of Pol II to promote cleavage and polyadenylation of the nascent pre-mRNA (62). However, Pol III lacks a homologous CTD and it is unclear whether Pol III transcripts could be similarly polyadenylated at their transcribed locus and, if so, how this might be stimulated following infection. To determine whether the CPSF complex could be recruited to transcriptionally active B2 SINE genes, we measured the occupancy of CPSF30 in mock- and MHV68-infected murine NIH3T3 fibroblasts by ChIP-seq. This revealed infection-induced CPSF30 recruitment to Polr3A-occupied, highly expressed B2 SINE loci (n=107, RPKM≥5000, from (47)) (**Fig. 4 A and B**). While the overall CPSF30 ChIP-seq signal was much lower than the Polr3A ChIP-seq signal, we nonetheless observed discernable CPSF30-occupied peaks at these highly expressed B2 SINE genes. There was also a strong correlation between Polr3A and CPSF30 ChIP-seq coverage at these loci, suggesting that high Polr3A occupancy may contribute to CPSF30 recruitment (**Fig. 4C**). CPSF30 occupancy and its correlation with Polr3A binding was still detectable, but at reduced levels when averaged across all Polr3A occupied B2 SINEs expressed during infection (n=3434, RPKM≥5) (**Fig. S3 A and B**).

**Figure 4.**
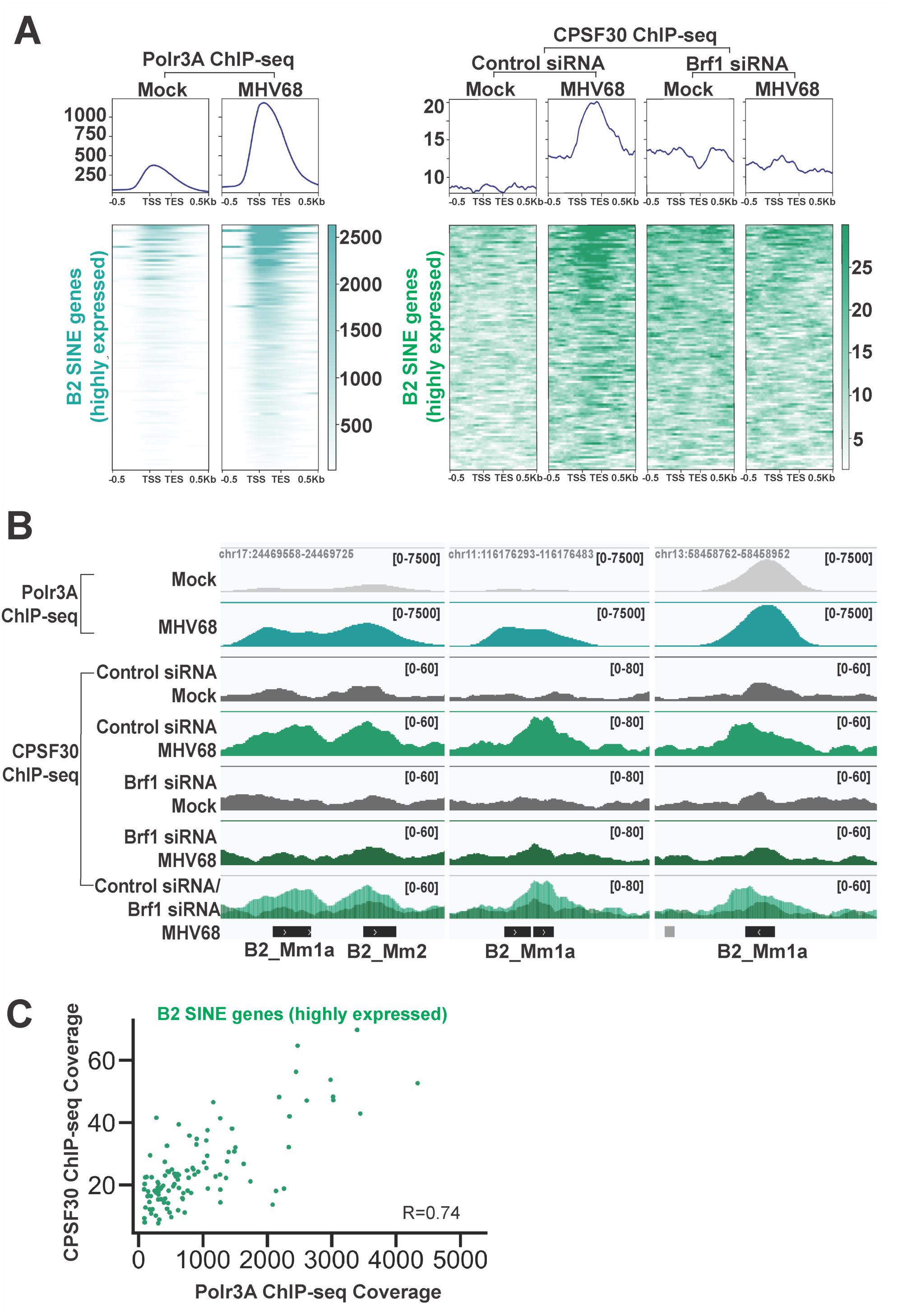
CPSF30 is recruited to B2 SINE genes during MHV68 infection (A) NIH3T3 cells transfected with control non-targeting or Brf1 targeting siRNAs were mock-treated or infected with MHV68 at an MOI of 5. At 24hpi, cells were harvested for ChIP-seq. The top panels are metagene plots displaying Polr3A or CPSF30 ChIP-seq signal across B2 SINE genes (n=107) with detectable Polr3A peaks (from Fig.1) and high expression levels (RPKM≥5000) (47). The Alignment files were averaged from two replicates. ChIP-seq signal was plotted as a histogram with 10 bp bins for −500 to +500bp around the TSS with 30 bins/gene. In the bottom panel, each row of the heat map corresponds to the CPSF30 ChIP-seq signal for each B2 SINE gene from the metaplot above. The heat maps are ranked by either Polr3A (left) or CPSF30 (right) ChIP-seq coverage values. (B) CPSF30 ChIP-seq coverage across select B2 SINE genes from NIH3T3 cells transfected with control non-targeting or Brf1 targeting siRNAs that were mock-treated or infected with MHV68. Alignment files were averaged from two replicates and visualized in IGV (96). Y-axis maximum and minimum values are within brackets. (C) Polr3A ChIP-seq signals associated with Polr3A occupied peaks at B2 SINE genes (n=107) with detectable Polr3A peaks (from Fig.1) and high expression levels (RPKM≥5000) (47) were plotted against CPSF30-ChIP-seq signals at the same loci. The correlation (R) between ChIP-seq signals is denoted on the graphs and was calculated using the Pearson correlation coefficient.

To assess whether Pol III recruitment was required for CPSF30 occupancy at B2 SINE loci, we performed CPSF30 ChIP-seq following siRNA-mediated depletion of Brf1, an essential subunit of the Pol III transcription factor TFIIIB that is required to recruit Pol III to type II SINE promoters (63). CPSF30 no longer occupied highly expressed B2 SINE loci in MHV68 infected cells lacking Brf1, indicating that Pol III occupancy is essential for virus-induced recruitment of CPSF30 to B2 SINE loci (**Fig. 4 A and B**). We confirmed that this loss of CPSF30 ChIP-seq signal was not due to a change in the abundance of CPSF30, which remained similarly expressed in the presence or absence of Brf1 (**Fig. S3C**).

### B2 SINE ncRNAs interact with the complete mRNA cleavage and polyadenylation complex during MHV68 infection

Our finding that CPSF30 was recruited to B2 SINE loci suggests that infection stimulates co-transcriptional polyadenylation of B2 SINE ncRNAs. To evaluate this further, we identified the proteins bound to B2 SINE ncRNAs in mock and MHV68 infected murine NIH3T3 fibroblasts using comprehensive identification of RNA-binding proteins (ChIRP) with B2 SINE-specific antisense oligonucleotides (**Fig. 5A**) (45). We probed the ChIRP protein eluates for components of the CPSF complex by western blot, and we were able to detect interactions between B2 SINE ncRNAs and several components of the CPSF complex (**Fig. 5B**). Additionally, ChIRP coupled to mass spectrometry (ChIRP-MS) recovered the complete CPSF complex, as well as components of the mRNA cleavage stimulatory complex (CSTF64 and CSTF77), and several accessory proteins involved in the regulation of CPSF-facilitated polyadenylation (PP1α, PP1β, and PABPN) (**Fig. 5C and Dataset S2**). The presence of the entire CPSF complex following B2 SINE ncRNA purification supports a transcription-coupled mechanism of B2 SINE ncRNA polyadenylation during MHV68 infection.

**Figure 5.**
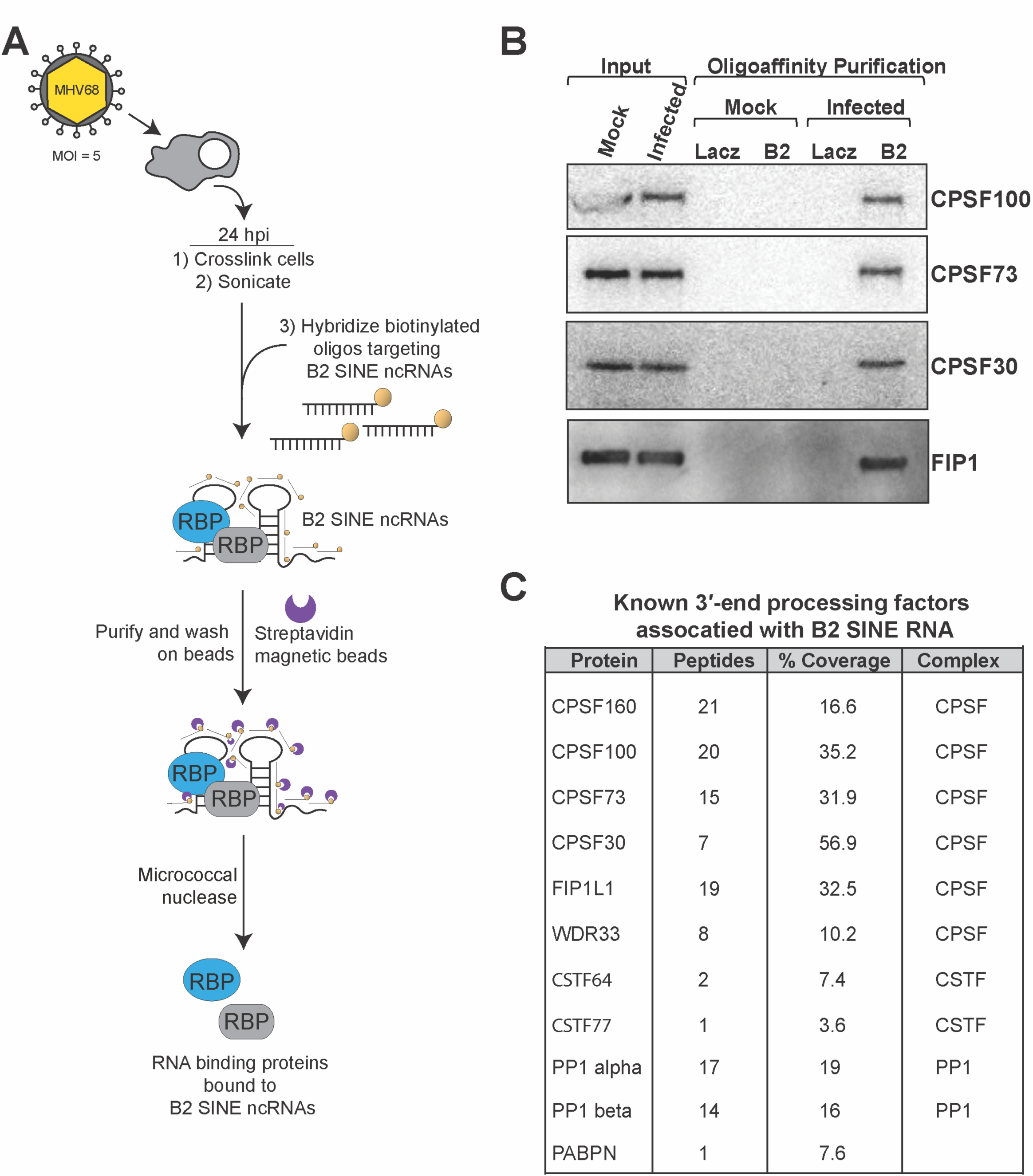
B2 SINE RNAs interact with mRNA 3**′**-end processing factors during MHV68 infection (A) Schematic of comprehensive identification of RNA-binding proteins (ChIRP) protocol to detect B2 SINE ncRNA-protein interactions during MHV68 infection. (B) NIH3T3 cells were mock-treated or infected with MHV68 at an MOI of 5. At 24 hpi, ChIRP was performed using biotinylated B2 SINE oligos. Antisense LacZ oligos were used as a negative control. Protein was extracted from ChIRP samples and analyzed by Western blotting with antibodies against CPSF100, CPSF73, CPSF30, and FIP1. (C) Total protein samples from panel (B) were also subjected to mass-spectrometry. The table shows peptide counts and percent coverage detected in samples associated with mRNA CPSF machinery and other 3′ processing factors.

### CPSF30 is also recruited to tRNA and Alu loci

We next investigated whether MHV68 infection induced CPSF30 recruitment to any other Pol III transcribed loci. While we did not detect CPSF30 at genes with type I (5S rRNA) and type III (U6 RNA, 7SK RNA, and Y RNA) promoters (**Fig. S4 A and B**), we did detect CPSF30 at Polr3A-occupied tRNA (n=295) genes during infection **(Fig. 6 A and B**). Like B2 SINE genes, CPSF30 ChIP-seq coverage at tRNAs was both MHV68 infection-induced and dependent on Brf1. However, we also observed a modest CPSF30 ChIP-seq signal at tRNAs in mock-infected cells (**Fig. 6A**). Furthermore, there was a stronger correlation between Polr3A occupancy levels and CPSF30 occupancy at tRNA genes compared to B2 SINEs during infection (**Fig. 6C and Fig. S3B**).

**Figure 6.**
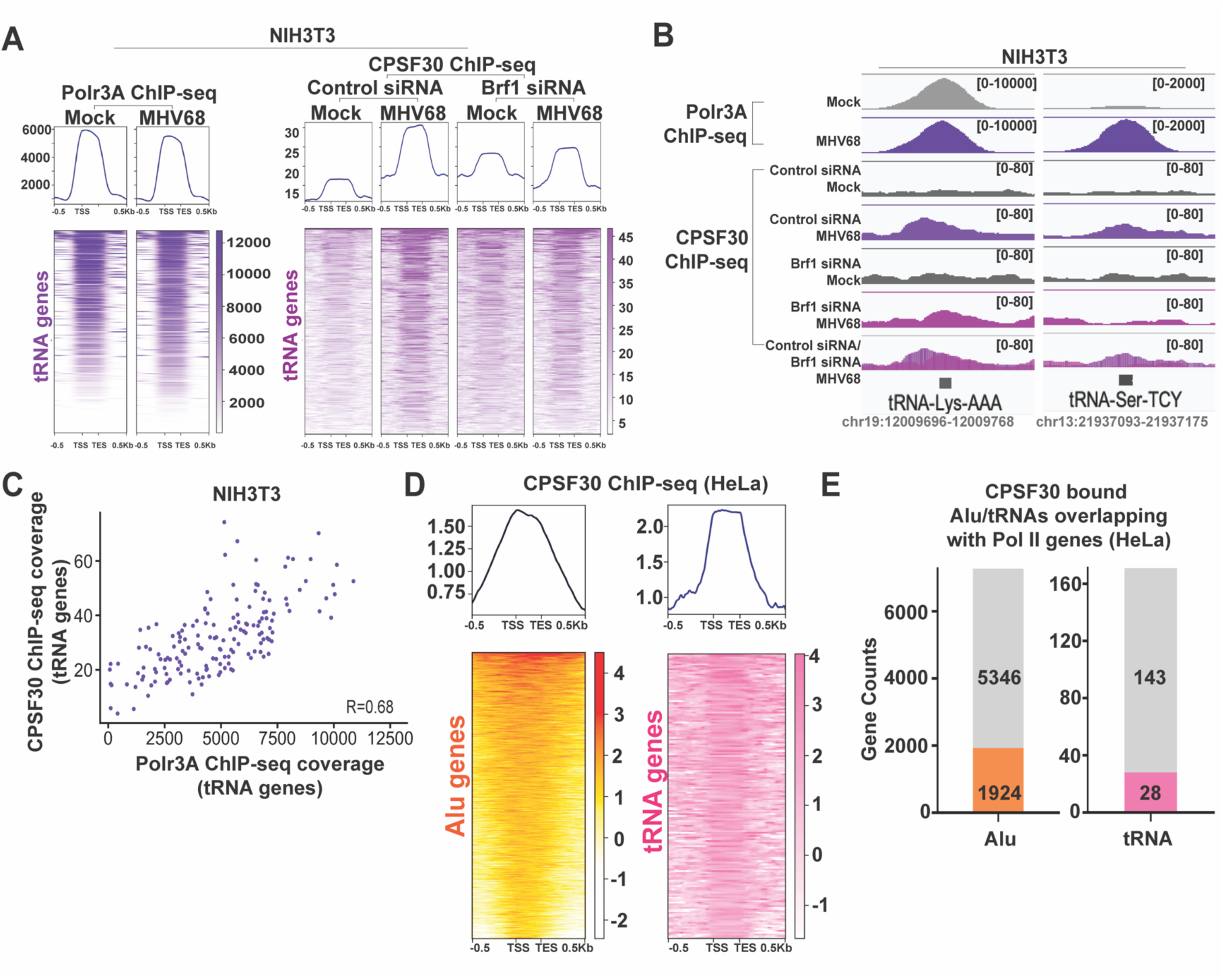
CPSF30 is recruited to tRNA and Alu genes (A) NIH3T3 cells transfected with control non-targeting or Brf1 targeting siRNAs were mock-treated or infected with MHV68 at an MOI of 5. At 24hpi, cells were harvested for ChIP-seq. The top panels are metagene plots displaying Polr3A or CPSF30 ChIP-seq signal across tRNA (n=295) genes with detectable Polr3A occupied peaks during infection (from Fig.1). Alignment files were averaged from two replicates. ChIP-seq signal was plotted as a histogram with 10 bp bins for −500 to +500bp around the TSS with 30bins/gene. In the bottom panel, each row of the heat map corresponds to the ChIP-seq signal for each tRNA gene from the metaplot above. Heat maps are ranked by either Polr3A (left) or CPSF30 (right) ChIP-seq coverage values. (B) CPSF30 ChIP-seq coverage across select tRNA genes from NIH3T3 cells transfected with control non-targeting or Brf1 targeting siRNAs and were mock-treated or infected with MHV68 at an MOI of 5. Alignment files were averaged from two replicates and visualized in IGV (96). Y-axis maximum and minimum values are within brackets. (C) Polr3A ChIP-seq signals associated with Polr3A occupied peaks at tRNA genes during infection (from Fig. 1) were plotted against CPSF30-ChIP-seq signals. The correlation (R) between ChIP-seq signals is denoted on the graphs and was calculated using the Pearson correlation coefficient. (D) The top panels are metagene plots displaying ChIP-seq signals at Alu (n=7270) or tRNA (n=171) genes that contain CPSF30 ChIP-seq peaks, called using the MACS2 (52) algorithm, from previously published CPSF30 ChIP-seq datasets generated from HeLa cells transfected with control antisense morpholino oligonucleotides (64). Alignment files were averaged from two replicates. ChIP-seq signal was plotted as a histogram with 10 bp bins for −500 to +500bp around the TSS with 30 bins/gene. In the bottom panel, each row of the heat map corresponds to ChIP-seq signal for each Alu or tRNA gene from the metaplot above. Heat maps are ranked by CPSF30 ChIP-seq coverage values. (E) Bar plots show the number of CPSF30 occupied Alu (left – orange) or tRNA (right – pink) genes from panel (D) that are embedded within Pol III transcribed genes. The number of Alu/tRNA genes that are bound by CPSF30 but no embedded within Pol II genes are shown in grey.

Finally, we asked whether CPSF30 recruitment to tRNA and SINE genes was a shared feature among mammalian genomes. Alu elements, which are evolutionarily distinct from B2 SINEs, are the largest subclass of infection-induced human SINEs. They are derived from the 7SL RNA and do not canonically contain genomically encoded PAS sequences. We analyzed CPSF30 occupancy at Alu (n=7270) and tRNA (n=171) genes from a previously published CPSF30 ChIP-seq dataset in human HeLa cells (64). Strikingly, when we used MACS2 (52) to peak call on the dataset, we observed prominent CPSF30 peaks at these genes, even in the absence of infection (**Fig. 6D**). Previous studies have shown that human Alu sequences contain sites that give rise to polyadenylation signals when embedded within Pol II transcribed protein-coding genes (12). This led us to investigate whether the Alu and tRNA genes bound by CPSF30 in HeLa cells were preferentially located within Pol II-transcribed genes. Our analysis revealed that only ∼26% of Alu genes and ∼16% of tRNA genes bound by CPSF30 overlapped with Pol II-transcribed genes (**Fig. 6E**), suggesting that localization within Pol II genes was not a major determinant of CPSF30 occupancy at these loci. Overall, CPSF30 occupancy at select Pol III loci is a feature of both murine and human genomes, suggesting new roles for the mRNA polyadenylation machinery in regulating and/or processing tRNA and noncoding retrotransposons.

## DISCUSSION

The ncRNAs derived from the thousands of SINE loci have significant gene regulatory, mutagenic, and inflammatory potential, and thus their expression is tightly regulated. Infection with the herpesvirus MHV68 is a notable example in which the constraints preventing B2 SINE ncRNA accumulation are somehow lifted. Here, we reveal that the dramatic accumulation of these retrotransposon ncRNAs during infection is due to a combination of Pol III transcriptional induction and recruitment of the CPSF machinery to B2 SINE loci, leading to Pol III transcription-associated polyadenylation of B2 SINE ncRNAs (**Fig. 7**). While Pol III occupies many loci even in uninfected cells, CPSF recruitment to these genes preferentially occurs in the context of infection, suggesting an inducible coupling of Pol III transcription and polyadenylation. An additional implication is that CPSF recruitment to SINE loci may, under some circumstances, enhance the potential for the polyadenylation-dependent retrotransposition of B2 SINEs, which is relevant considering that many SINE-inducing DNA viruses are oncogenic.

**Figure 7.**
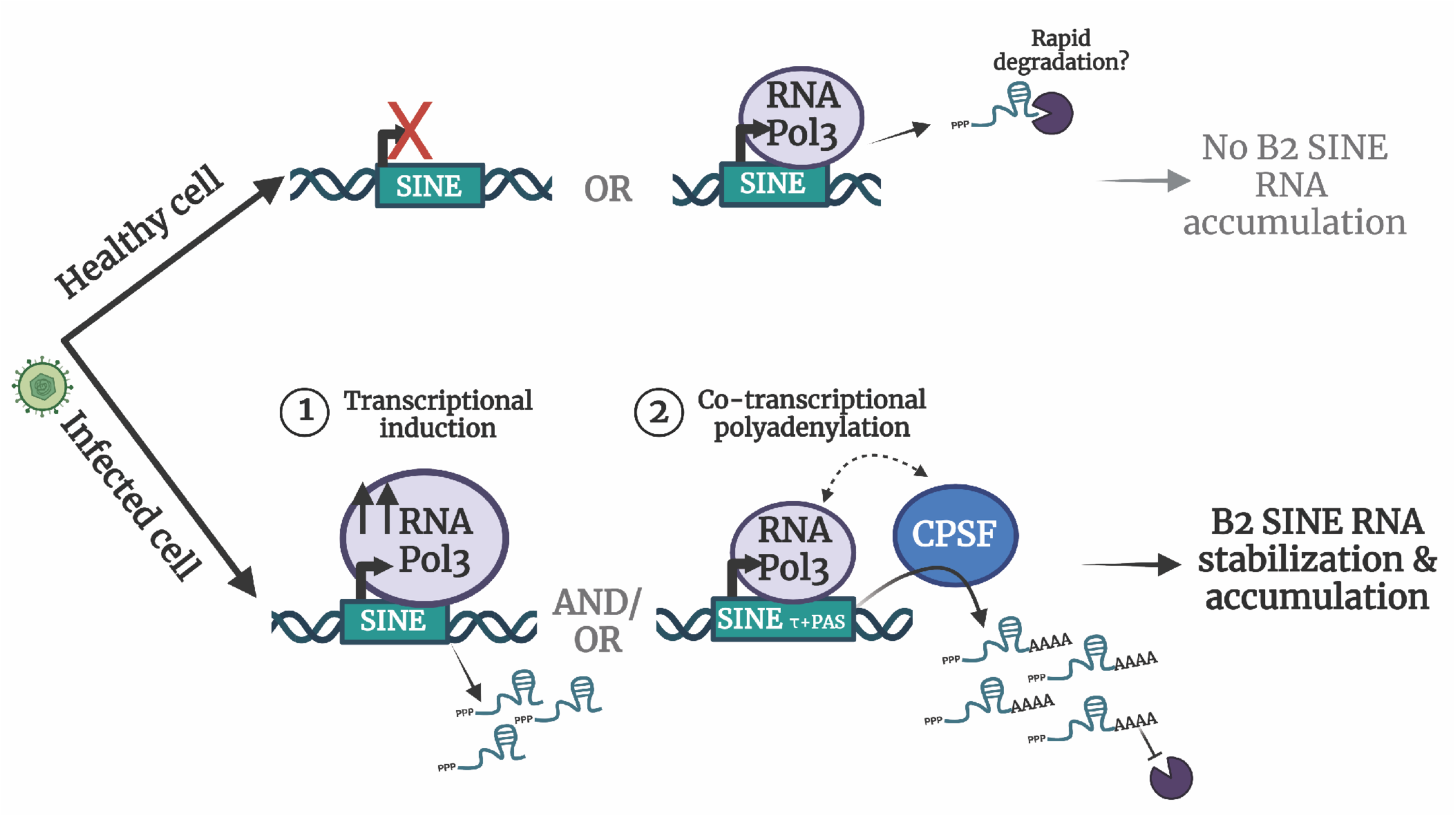
Model of B2 SINE induction and polyadenylation during MHV68 infection. Schematic shows possible models for B2 SINE ncRNA induction during MHV68 infection. The top panel shows B2 SINE loci in healthy cells, which are repressed or occupied by Pol III, with B2 SINE ncRNAs likely being rapidly degraded. The bottom panel shows models for transcriptional induction at B2 SINE genes (1) and co-transcriptional polyadenylation of B2 SINE ncRNAs via CPSF activity (2), both leading to the accumulation of SINE ncRNAs in virus-infected cells.

Deep learning models have emerged as remarkably accurate predictors of gene regulatory features. Recent examples include predicting mRNA abundance (65), transcription factor binding (66–68), Pol II pausing (69), and Pol III binding to tRNA genes (70), as well as cis-regulatory motifs for mRNA poly(A) site selection, strength, and cleavage (71–74). As we demonstrate here with SAMBAR-Net, convolutional neural networks can also overcome challenges associated with discriminative analysis of highly repetitive, hyperabundant, and degenerate sequences like SINEs. The ∼350,000 B2 SINEs in the mouse genome all contain the same internal type II A/B box promoter. Yet, only a subset of those with intact promoters produce detectable B2 SINE ncRNA in response to infection (47). While MEME identified long stretches of DNA sequence that discriminate B2 SINE ncRNA abundance during MHV68 infection, SAMBAR-Net provided base pair resolution of key nucleotides within the polyadenylation-promoting sequence motif. Unlike the more closely related B2_Mm1a, B2_Mm1t, and B2_Mm2 SINE families, B3 and B3A SINE genes are known to be more divergent and do not contain intact τ motifs, which may underlie their low expression in infected cells (19). We envision several possible future applications of CNNs that could extend our understanding of Pol III regulation in healthy versus stressed cells. This includes whether any of the CNN-based findings regarding mRNA poly(A) regulation apply to other SINEs or tRNAs, as well as whether surrounding sequence features outside of the SINE gene body dictate Pol III binding. With respect to this latter possibility, our ChIP-seq data show that there is specificity in Polr3A occupancy at B2 SINE loci, even in unperturbed cells, suggesting that chromatin features may also influence B2 SINE locus accessibility (75). It is intriguing that tRNA genes and human Alu elements also recruit CPSF30 even though they lack the PAS and τ motif required by B2 SINEs, as this indicates that features directing the polyadenylation machinery to Pol III loci have not yet been exhaustively defined.

Polyadenylation is known to stabilize B2 SINE ncRNAs (21), which explains how its induction increases the abundance of these normally unstable Pol III transcripts. Beyond the canonical T-tract termination signal in the non-template strand of Pol III transcripts, a subset of B2 SINEs (Class T^+^) contain an internal PAS together with a τ motif, which is required for efficient cleavage and polyadenylation by the CPSF complex (17–21, 61). For mRNA polyadenylation, current models suggest that the CPSF complex is recruited to the 5′-end of the transcript, possibly coordinately by TFIID and the CTD of Pol II (62, 76). Given that Pol III lacks an analogous CTD, one hypothesis is that PAS-containing B2 SINE ncRNAs are polyadenylated in the nucleoplasm after canonical Pol III termination rather than the machinery being recruited directly to the sites of transcription. While we cannot exclude this possibility, our observation that CPSF machinery is recruited to B2 SINE loci in response to viral infection supports the hypothesis that polyadenylation of B2 SINE ncRNA can occur co-transcriptionally during infection (**Fig. 7**). The idea that MHV68-induced polyadenylation of B2 SINE ncRNAs is tightly coupled to Pol III transcription is further supported by the observation that depletion of Brf1, an essential component of the Pol III transcription factor complex TFIIIB, prevents CPSF30 occupancy at B2 SINE loci. CPSF recruitment to Pol III loci is presumably at least partly an RNA-dependent process, as the B2 SINE ncRNA binds the entire mRNA-like CPSF complex. We hypothesize that this recruitment and polyadenylation counteract the rapid degradation of B2 SINE ncRNA that occurs when it is transcribed in uninfected conditions (**Fig. 7**).

How the CPSF complex is recruited to Pol III loci, particularly in an infection-stimulated manner, is a key open question. One hypothesis is that MHV68 infection induces remodeling of the Pol III transcription pre-initiation complex, for example, causing the recruitment of a factor(s) with CPSF binding sites or altering the regulation of Pol III itself in a manner that promotes CPSF binding. Our data suggest that type II, but not type I or II Pol III initiation complexes, are associated with infection-induced CPSF30 recruitment, suggesting a possible promoter-specific recruitment mechanism. Indeed, a variety of viruses including herpesviruses can alter Pol II subunit abundance and modification via mechanisms with parallels to cell stress pathways (77). Additionally, Pol II occupancy has been reported at or near Pol III transcribed genes (78–84), particularly tRNA genes, which could enable a Pol II-linked handoff mechanism at those sites. However, our observation that CPSF30 is more prominently associated with Pol III loci than with its canonical Pol II loci suggests a unique mechanism of recruitment. Another possibility is that the presence of the τ motif boosts the ability of an infection-modified Pol III complex to recruit CPSF30. If so, the high level of Pol III occupancy at tRNA genes may underlie their ability to recruit CPSF30 in a τ motif independent manner, whereas B2 SINE loci, which even in infected cells display much lower levels of Pol III occupancy, may require the assistance of a τ motif.

The functional outcomes of polyadenylation-induced Pol III transcript abundance during viral infection are likely to be varied and context-specific. For example, the induction of polyadenylated B2 SINE ncRNAs by MHV68 can trigger host responses, including activation of the antiviral NF-kB signaling pathway and altering mRNA export in a transcript-specific manner through RNA-RNA interactions (45, 47). MHV68 also co-opts Pol III to transcribe the viral TMER loci that are processed into miRNAs involved in regulating infection and virulence *in vivo* (85–87). However, there are several additional DNA viruses that lack Pol III genes on their genomes (e.g., HSV-1) whose infections lead to modulation of Pol III transcription and CPSF activity on both the host and viral genomes (49, 88). This suggests a much broader role for Pol III transcription and polyadenylation of ncRNAs during viral infection. Deciphering how viruses modulate Pol III transcription and RNA processing should also be relevant for other cellular stresses, as well as continuing to uncover mechanisms governing fundamental gene expression.

## METHODS

### Cell lines

NIH 3T3 (ATCC CRL-1658) and NIH 3T12 (ATCC CCL-164) mouse fibroblast cell lines were maintained in Dulbecco’s modified Eagle’s medium (DMEM; Gibco) with 10% fetal bovine serum (FBS; VWR) and screened regularly for mycoplasma by PCR.

### siRNA nucleofections

Transfections of siRNAs into NIH 3T3 fibroblasts were completed as follows: Cells were maintained in DMEM with 10% FBS and grown to 90% confluence. Cells were removed and washed once with Dulbecco’s phosphate-buffered saline (DPBS) (Gibco). Nucleofections were done using the Neon Transfection System (Thermo Fisher). 2 × 10^6^ cells were resuspended in 100 μL of buffer R, to which control non-targeting, Brf1, CPSF30, CPSF73, or Nudt21 pools of siRNA (ON-TARGETplus SMARTpool siRNA – Horizon Discovery) were added to a final concentration of 200nM. This was loaded into a Neon 100-μL pipette tip and Neon tube with 3 mL of buffer E2 with electroporation parameters set to 1,300 V, 20 ms, 2 pulses. Following electroporation, cells were plated in 10 mL of DMEM with 10% FBS in 10 cm TC-treated plates and were incubated at 37°C for 24 hours.

### Virus preparations and infections

MHV68 was amplified in NIH 3T12 fibroblast cells, and the viral 50% tissue culture infective dose (TCID50) was measured on NIH 3T3 fibroblasts by limiting dilution. NIH 3T3 fibroblasts were infected at the indicated multiplicity of infection (MOI) by adding the required volume of the virus to cells in 5 mL of serum-free DMEM in 10-cm TC-treated plates. Infection was allowed to proceed for one hour to allow for viral entry, followed by removal of the virus-containing media and replacement with fresh DMEM with 10% FBS. Cells were harvested 24 hours post-infection.

### Chromatin immunoprecipitation (ChIP)

NIH3T3 mouse fibroblast cells plated in a 10-cm TC-treated dish were cross-linked in 1% formaldehyde in DPBS (Gibco) for 5 minutes at room temperature, quenched in 0.125 M glycine, and washed twice with DPBS (Gibco). Cross-linked cells were harvested and lysed with 1 ml ChIP lysis buffer (50 mM HEPES [pH 7.9], 140 mM NaCl, 1 mM EDTA, 10% glycerol, 0.5% NP-40, 0.25% Triton X-100) by rotation for 10 min. Nuclei were collected by centrifugation at 1,700 × *g* for 5 min at 4°C, washed once with ChIP shearing buffer (50 mM Tris-Cl, pH 7.5, 10 mM EDTA, 0.1% SDS), and then resuspended in 1 ml of ChIP shearing buffer. Chromatin was then sheared for 5 min using a Covaris S220 focused ultrasonicator at 140 power, a 5% duty cycle, and 200 bursts/cycle. Chromatin was spun at 15,000 × *g* for 5 min at 4°C, and the pellet was discarded. 40 μg of chromatin was incubated with 10 μg rabbit polyclonal anti-POLR3A (Abcam ab96328), anti-CPSF30 (Proteintech 15023-1-AP), or rabbit IgG (Abcam ab37415) overnight. 25 μl of mixed protein A and G Dynabeads (Thermo Fisher) were added, and the tubes were rotated for 2 hours at 4°C. Dynabeads were then washed with low-salt immune complex (20 mM Tris [pH 8.0], 1% Triton X-100, 2 mM EDTA, 150 mM NaCl, 0.1% SDS), high-salt immune complex (20 mM Tris [pH 8.0], 1% Triton X-100, 2 mM EDTA, 500 mM NaCl, 0.1% SDS), lithium chloride immune complex (10 mM Tris [pH 8.0], 0.25 M LiCl, 1% NP-40, 1% deoxycholic acid, 1 mM EDTA), and Tris-EDTA for 5 min each at 4°C with rotation. DNA was eluted from the beads using 100 μl of elution buffer (150 mM NaCl, 50 μg/ml proteinase K) and incubated at 50°C for 2 h and then at 65°C overnight. DNA was purified using an Oligo Clean and Concentrator kit (Zymo) and eluted in 50 μl of nuclease-free water. DNA was used for generating libraries for ChIP-seq.

### ChIP sequencing and data analysis

Libraries for ChIP-seq were prepared using the NEBNext Ultra II DNA Library Prep Kit for Illumina (New England Biolabs) with NEBNext Multiplex Oligos for Illumina (Unique Dual Index UMI Adaptors DNA Set 1). Equal amounts of sample eluted DNA and spike-in DNA (yeast spike-in DNA – Cell Signaling) were used within each experiment and amplified for 3-10 cycles based on NEB recommendations and sequenced on a NovaSeq X 10B (Illumina) with 150 paired-end reads. Sequencing quality was assessed with FastQC, and all sequencing files were processed using HTStream with default settings for paired-end reads, including trimming adapter sequences, PhiX screening, “N” nucleotide trimming, and removing low-quality reads. The mm10, yeast (GCA_000146045.2), and MHV68 (U97553.2) genomes were manually indexed using the bowtie2-build command. Processed sequencing files were then separately aligned to the three genomes with bowtie2(89) using the following parameters: --maxins 1500, -k 25, --no-mixed, and --no-discordant to allow for multiple mapping locations per read. Allo (90) algorithm was used to redistribute multimapping reads, including those that map to highly repetitive Pol III-transcribed genes. The resulting bam files were further filtered to discard any unmapped reads and supplementary alignments (samtools -F 2048), ensure reads are properly paired (samtools -f 2), and blacklist regions on the mm10 genome (bedtools intersect -v)(91, 92). Peak calling was performed with MACS2 (52) on Polr3A ChIP-seq data with input normalization and default settings except for setting --min-length to 150 to capture small Pol III-transcribed genes. Bedtools intersect -u was performed on the mm10 RepeatMasker (93) file and called peaks to identify the precise loci that were bound by Polr3A. Consensus Polr3A-bound genes were then identified using bedtools intersect between replicates with the -f 1, -r, and -u options. All consensus Polr3A-bound genes in the mock and MHV68-infected conditions were concatenated into one file for differential binding analysis with DiffBind/DESeq2 (54, 94) using spike-in normalization scale factors proportional to the total number of spike-in reads. Spike-in normalization was used to compare across different sample conditions. Differentially bound regions were defined as FDR≤0.05 and a log_2_FC≥1.5. Polr3A coverage correlation plots with B2 SINE ncRNA or pre-tRNA expression levels were generated by using the bedtools map command to identify the max (-o max) coverage value from a bedGraph converted BigWig file for the specified genomic regions.

### ChIP-seq visualization

Coverage profiles generated for all ChIP-seq datasets were produced using DeepTools (95). For Polr3A and CPSF30 ChIP-seq data in NIH3T3 cells, BigWig files were first generated with the bamCoverage command for each bam file from each sample mapped to the mm10 genome with the following parameters --binSize 10, --smoothLength 30, --extendReads, --centerReads, -- normalizeUsing None, and --scaleFactor with their respective spike-in normalized scale factor (reciprocal of the ones used in DiffBind). CPSF30 ChIP-seq data in HeLa cells were similarly processed but were instead log_2_ input-normalized with the bamCompare command with the following changes in parameters: --scaleFactorsMethod None, --normalizeUsing RPKM, and – operation log2. Average BigWig file was then generated for each biological condition with the bigwigAverage command containing the -bs 10 option. The calculation for heatmap of Polr3A and CPSF30 coverage profiles was generated using the computeMatrix scale-regions command with the average BigWig files (-S), the mm10 blacklist option (-bl) for redundancy, --skipZeros, -- missingDataAsZero, and the following options for visualization purposes: -- beforeRegionStartLength 500, --regionBodyLength 300, --afterRegionStartLength 500. For Polr3A ChIP-seq data in NIH3T3 cells and CPSF30 ChIP-seq data in HeLa cells, the lists of regions were generated from Pol III-transcribed RepeatMasker genes that overlap with called peaks. For CPSF30 ChIP-seq data in NIH3T3 cells, coverage profiles were generated for all (RPKM≥5) or highly expressed (RPKM≥5000) B2 SINE and tRNA genes in RepeatMasker(93) that overlap with called peaks. Heatmaps and metaplots were then generated with the plotHeatmap command. All genome tracks shown are from the Integrative Genome Viewer(96) with the DeepTools-produced averaged BigWig files. ChIP-seq data have been deposited in NCBI’s Gene Expression Omnibus and are accessible through GEO accession number **GSE288677**.

### MEME analysis

MEME (60) was used in discriminative mode to identify sequence features that discriminated between highly expressed and unexpressed B2 SINE loci as defined by having an RPKM≥10 and RPKM=0, respectively. The input data were taken from a previously generated SINE-seq dataset (47), and DNA sequences were extracted from the mm10 genome FASTA file. All settings in MEME were set to default.

### SAMBAR-Net architecture

SAMBAR-Net is a binary CNN model implemented using PyTorch (97) to discriminate between expressed (RPKM≥10) from unexpressed (RPKM=0) B2 SINE genes in NIH3T3 cells using a one-hot encoded format as its input. The input data was obtained from our previously generated SINE-seq dataset (47) and labeled based on each B2 SINE gene’s expression level. The B2_Mm1a, B2_Mm1t, and B2_Mm2 B2 SINE families were focused on, as expressed B3 and B3A SINEs are severely underrepresented in the SINE-seq dataset. The length of each B2 SINE sequence began at the TSS and extended 300 base pairs downstream to accommodate for the average length of a B2 SINE gene. Due to the repetitive nature of B2 SINE genes within a family, a subset of the training and test sets could have highly similar sequences, which could cause the model to overfit quickly. To prevent overfitting, a simple DenseNet-like (55) architecture was developed with only three convolutional layers and two dense blocks – one between each convolutional layer – to reduce the number of parameters to train. Each convolutional layer contained 128 filters with a width of 10 base pairs. Batch normalization, ReLu activation, maxpooling (kernel size=5, stride=1), and dropout (probability=0.3) layers were applied to each convolutional layer. Each dense block contained three convolutional layers, each with 128 filters with a width of 3 base pairs, padding of 2, and a growth rate of 32, in which each layer is directly connected to every other layer in a block. Each convolutional layer in a dense block had batch normalization and ReLu activation layers applied.

### CNN training and evaluation

The CNN was trained using an 80-10-10 split of the B2 SINE expression dataset with stratification for each family to retain similar gene counts of each B2 SINE family in the training, validation, and testing datasets. The model was trained for 1500 epochs, and the model at the epoch producing the lowest validation loss was saved and analyzed on the testing dataset. The training was done in batch sizes of 64 with the Adam optimizer (learning rate = 1e-7), and the model was evaluated on the held-out test set by measuring precision, recall, F1 score, and AUC. AUROC curves were plotted for each of the three B2 SINE families analyzed to determine model performance on each family independently. TF-MoDISco (56) was used to identify the DNA sequence features that contribute to the model’s discrimination between expressed and unexpressed B2 SINE loci for each family. All settings used in TF-MoDISco were set to defaults, except that each B2 SINE family contained unique motif length discovery settings when being analyzed.

Position frequency matrix (PFM) was extracted from TF-MoDISco and converted to a position weight matrix (PWM) by normalizing the frequencies to probabilities and dividing each probability in the matrix by the background frequency of nucleotides for the mm10 genome (A=0.29, C=0.21, G=0.21, T=0.29). PWMScan (59) was used to calculate PWMScores for expressed B2 SINEs as defined in the SINE-seq dataset (47). PWMScores reflect the strength of prediction for the binding site identified through TF-MoDISco. All settings were set to default when using PWMScan. Expressed B2 SINEs were then divided into quartiles based on their PWMScores.

### Northern blotting

Total RNA was extracted from cells using TRIzol reagent (Invitrogen). Northern blotting for small RNAs was carried out as previously described with some modifications (98). 10 μg of total RNA was separated by electrophoresis on 8% TBE-Urea gels and transferred onto a Hybond N^+^ membrane (Amersham). If RnaseH digestions were performed, 10 µg of total RNA was combined with 500 pmol of oligo(dT) primer in a 25 µl reaction, incubating at 65°C for 8 min, then adding 1 U of RnaseH (New England Biolabs), RnaseH buffer, and 40 U of Rnasin (Promega). Reactions were incubated at 37°C for 30 min, then terminated by adding 1 µl of 0.5 M EDTA (pH 8.0) and precipitating the RNA in 1ml of 100% ethanol before gel electrophoresis. Membranes were cross-linked at 2400 J/m^2^ (UV Crosslinker, VWR). B2 SINE ncRNA was detected using the following digoxigenin-labeled (DIG) end-labeled oligo: GATGGTTGTGAGCCACCATGTGGTTGCTGGCA. Oligos were end-labeled using a DIG Oligonucleotide Tailing Kit (Roche). Gels were stained with 1 μg/ml Sybr Safe (Invitrogen) to detect 5.8S and 5S rRNAs as loading controls.

### Primer extension

Total RNA was extracted from cells using TRIzol reagent (Invitrogen). Primer extension was performed on 15 μg of total RNA using 5′-fluorescein-labeled oligonucleotides specific to B2 SINE RNA (56-FAM/tacactgtagctgtcttcagaca) and 7SK RNA (/56-FAM/tgagcttgtttggaggttct). RNA was ethanol precipitated in 1 mL 100% EtOH, washed in 70% ethanol, and pelleted at 21,130 × *g* and 4°C for 10 min. Pellets were resuspended in 18 μL of 1X SuperScript III reverse transcriptase reaction buffer (SSIII-RT; Thermo Fisher) containing 1 μL of each 5′-fluorescein-labeled primer (10 pmol/μL). Samples were heated to 80°C for 10 min, followed by annealing for 1 h at 56°C. Then, 30 μL of extension buffer (1X SSIII-RT buffer, 40U Rnasin Rmase Inhibitor [Promega] 2 mM DTT, 1 mM dNTP, 1,000 U of SSIII-RT) was added, and extension was carried out for 1 h at 42°C. Samples were precipitated in 100% ethanol for 20 min at −80°C, and then pellets were briefly air dried and resuspended in 20 μL 1× RNA loading dye (47.5% formamide, 0.01% SDS, 0.01% bromophenol blue, 0.005% xylene cyanol, and 0.5 mM EDTA). Then, each sample was run on an 8% urea-PAGE gel for 1 h at 250 V. Gels were imaged on a Chemidoc imager (Bio-Rad) with fluorescein imaging capability. The relative induction of B2 SINE ncRNAs for each sample was measured as the ratio of the mean integrated intensity between 7SK RNA and B2 SINE ncRNA level using FIJI (99) was normalized to the mock-treated control.

### Western blotting

To prepare whole-cell lysates for evaluating protein expression of CPSF factors, mock-treated and MHV68-infected cells were washed with cold DPBS (Gibco) followed by lysis with radioimmunoprecipitation assay (RIPA) lysis buffer (50 mM Tris HCl, 150 mM NaCl, 1.0% [vol/vol] NP-40, 0.5% [wt/vol] sodium deoxycholate, 1.0 mM EDTA, and 0.1% [wt/vol] SDS, Roche cOmplete Mini EDTA-free protease inhibitor cocktail). Cell lysates were vortexed briefly, rotated at 4°C for 15 min, and then clarified by centrifugation at 21,000 × *g* in a tabletop centrifuge at 4°C for 10 min to remove debris. 30 μg of whole-cell lysate were resolved on 4% to 15% mini-PROTEAN TGX gels (Bio-Rad). Transfers to polyvinylidene difluoride (PVDF) membranes (Bio-Rad) were done with the Trans-Blot Turbo transfer system (Bio-Rad). Blots were incubated in 5% milk in TBS with 0.1% Tween 20 (TBS-T) to block, followed by incubation with primary antibodies against anti-Nudt21 (Proteintech 10322-2-AP, 1;1000), anti-Vinculin antibody (abcam ab91459, 1;5,000), anti-CPSF30 antibody (Proteintech 15023-1-1AP, 1;1000), anti-CPSF73 antibody (Bethyl Laboratories A301-090A, 1;1000 or abcam ab72295, 1;100), anti-CPSF100 antibody, or anti-FIP1 antibody. Washes were carried out with TBS-T. Blots were then incubated with HRP-conjugated secondary antibodies (Southern Biotechnology, 1:5,000). Washed blots were incubated with Clarity Western ECL substrate (Bio-Rad) for 5 min and visualized with a ChemiDoc imager (Bio-Rad).

### Comprehensive identification of RNA-binding proteins (ChIRP) and mass spectrometry

ChIRP was performed as previously described with minor modifications (45). Briefly, ∼ 100 million NIH3T3 cells were infected with MHV68 at an MOI 5. 24 hpi cells were cross-linked with 1.1% formaldehyde for 15 min at room temperature. Crosslinking was then quenched with 0.125 M glycine for 5 min. Cells were rinsed again with PBS, scraped into Falcon tubes, and pelleted at 1000 xg for 5 min. The cell pellet was resuspended in 3 mL nuclei lysis buffer (50 mM Tris-HCl [pH 7.0], 10 mM EDTA, 1% SDS, protease cocktail inhibitor [Roche], and RNase inhibitor [Fermentas]) and rotated for 10 min. at 4°C. Cells were dounced10 times with a B-type pestle and separated into three 1 mL aliquots for sonication. Sonication was performed using a Covaris-focused sonicator. After sonication, chromatin aliquots were combined, and 9 mL of hybridization buffer (750 mM NaCl, 1% SDS, 50 mM Tris 7.0, 1 mM EDTA, 15% Formamide, protease inhibitor cocktail, and RNAse inhibitor) was added. 50 pmol of five separate 3′-TEG biotinylated probes were added to the dilute chromatin and rotated overnight for 16 h. Streptavidin-magnetic C1 beads (Life Technologies) were washed three times in nuclei lysis buffer, blocked with 500 ng/μl yeast total RNA, and 1mg/ml BSA for 1 hr at room temperature, and washed three times again in nuclear lysis buffer before being resuspended in its original volume. One hundred microliters of washed/blocked C1 beads were added to the chromatin mixture and rotated for an additional 4 h at 37°C. Beads:biotin-probes:RNA:protein adducts were captured by magnets (Invitrogen) and washed five times with 10 mL wash buffer (2× SSC, 0.5% SDS). After the last wash, complexes were eluted by resuspending the beads in 500 μL G50 buffer (20 mM Tris-HCl, 300 mM NaCl, 2 mM EDTA, 0.2% SDS), The elution was split in half for isolation of protein and RNA. 50 μg/mL Proteinase K (Fermentas) was added to the fraction for nucleic acid isolation and incubated at 60°C for 1 h. The G50 buffer was separated from the beads, phenol-chloroform was extracted, and ethanol precipitated. RNA was extracted and analyzed by small RNA northern blotting. Protein was similarly isolated, except no Proteinase K was added. For mass-spectrometry, samples were provided to the University of California, San Francisco Mass Spectrometry facility for processing, trypsin digestion, and analysis by LC-MS.MS on a Thermo Scientific Velos Pro ion trap mass spectrometry system.

### DATA, MATERIALS, and SOFTWARE AVAILABILITY

Further information and requests for materials should be directed to the Lead Contact, Britt A. Glaunsinger.

All ChIP-seq data generated in this study have been deposited in NCBI’s Gene Expression Omnibus and accessible through GEO accession no**. GSE288677**. Previously generated SINE-seq data from mock- and MHV68-infected NIH3T3 cells analyzed in this study were obtained from GEO accession no. **GSE85518**. Previously generated DM-tRNA-seq data from mock- and MHV68-infected MC57G cells analyzed in this study were obtained from GEO accession no. **GSE142393**. Previously generated CPSF30 ChIP-seq data from HeLa cells were obtained from GEO accession no. **GSM5776370**. All ncRNA genome annotations are from RepeatMasker (93).

All code used to train and analyze SAMBAR-Net can be found at: https://github.com/UCB-GLab/SAMBAR-Net

## ACKNOWLEDGEMENTS

We thank all current and past members of the Glaunsinger, Lareau, and Coscoy Labs. We particularly thank Leah Gulyas, Chad Stein, Lidia Llacsahuanga, and Sam Rider for their careful reading of the manuscript. We thank Alma Burlingame and Kathy Li at the UCSF Mass Spectrometry Facility for performing the mass spectrometry and data analysis. Sequencing was performed through the Vincent J. Coates Genomics Sequencing Laboratory at UC Berkeley (QB3 Genomics, UC Berkeley, Berkeley, CA, RRID:SCR_022170). This work is funded by National Institutes of Health (NIH) R01CA136367 and R01AI122528 to B.A.G., who is also an investigator of the Howard Hughes Medical Institute.

## AUTHOR CONTRIBUTIONS

A.L., S.B.S., J.K., L.F.L., and B.A.G. designed research; A.L., S.B.S., X.M., P.S., and J.K. performed research; A.L., S.B.S., X.M., P.S., J.K., L.F.L., and B.A.G. analyzed data; A.L, S.B.S., and B.A.G. wrote the paper.

## DECLARATION OF INTERESTS

The authors declare no competing interests.

## SUPPLEMENTAL INFORMATION

Document S1: Figures S1-S4 Dataset S1 (separate file) Dataset S2 (separate file)

## SUPPLEMENTAL INFORMATION

**Figure S1.**
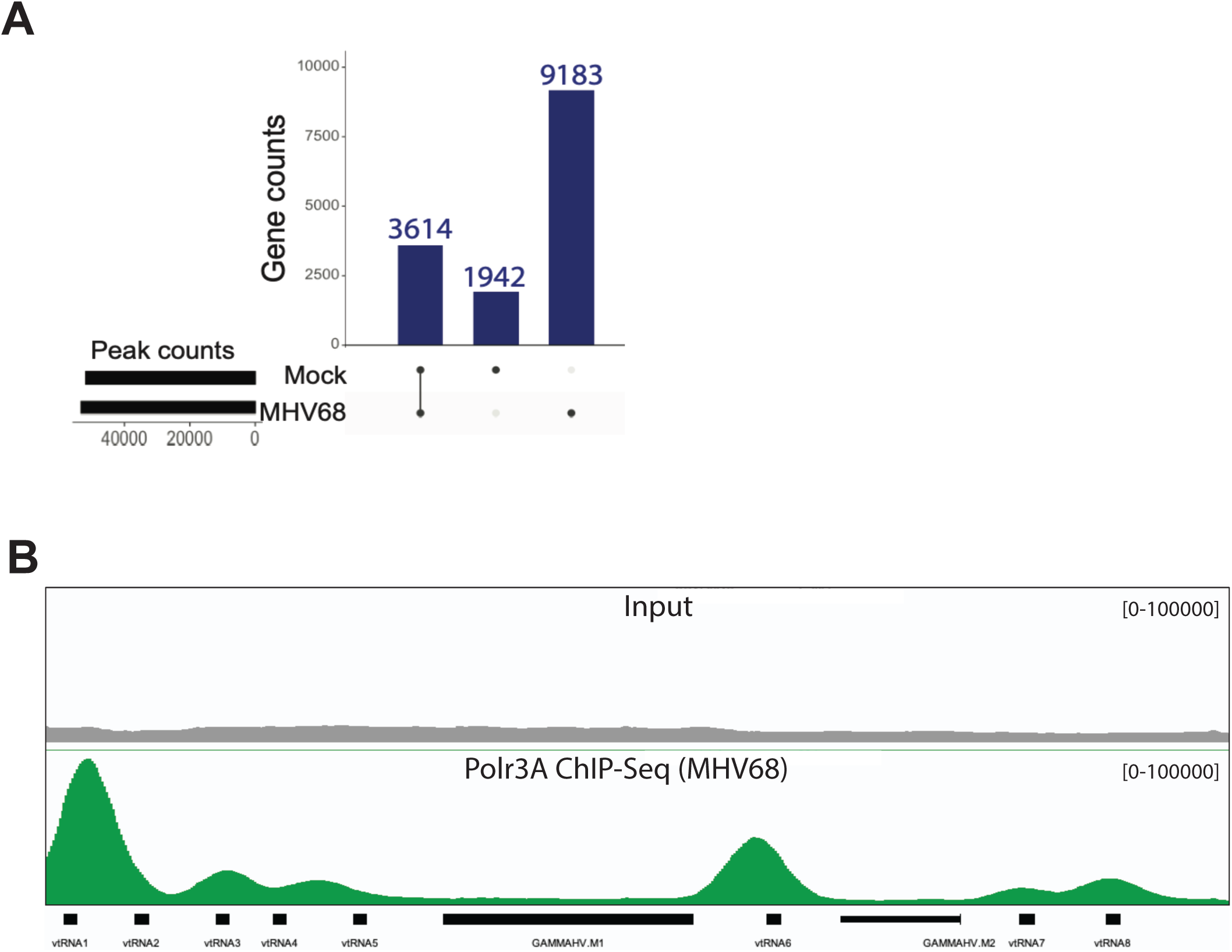
Recruitment of RNA Polymerase III to the MHV68 genome (A) MACS2 (1) peak calling algorithm was applied to Polr3A ChIP-seq data from mock-treated or MHV68-infected (MOI of 5 for 24 hours) NIH3T3 cells to determine the number of Polr3A bound peaks across the murine genome. Peak positions were then overlapped with RepeatMasker (2) gene annotations to determine which subset peaks overlapped with Pol III transcribed genes. The upset plot shows the number of peaks in each sample condition on the left and plotted on the right is the number of Pol III transcribed genes overlapping with these peaks that are unique to mock-treated or MHV68-infected samples or are present in both samples. (B) Polr3A ChIP-seq coverage across the MHV68 genome from MHV68-infected samples. Alignment files were averaged from two replicates and visualized in the Integrative Genome Viewer (3). Y-axis maximum and minimum values are within brackets. Viral TMER genes (vtRNA1-8) are shown below as a reference.

**Figure S2.**
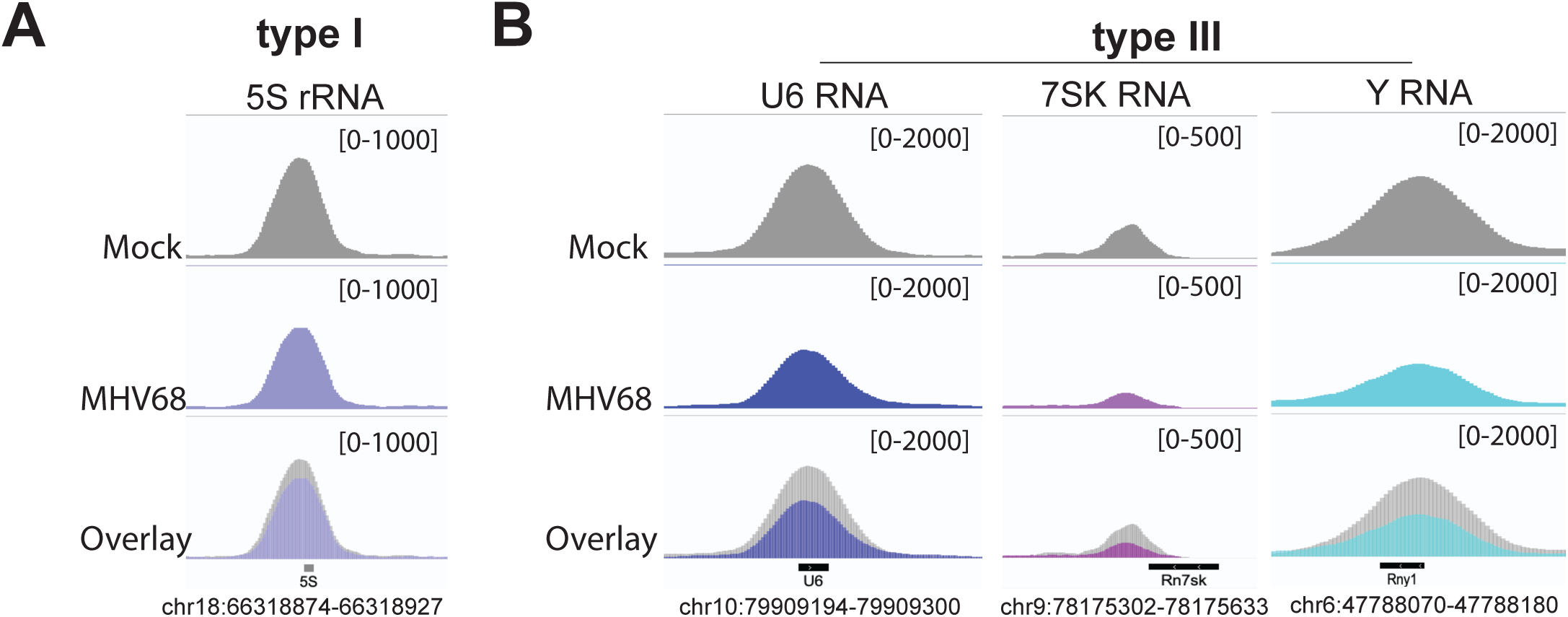
RNA Polymerase III binding to type I and type III promoters on the cellular genome during MHV68 infection Polr3A ChIP-seq coverage across select promoter type I (A) and type III (B) genes from mock-treated or MHV68-infected samples. Alignment files were averaged from two replicates and visualized in the Integrative Genome Viewer (3). Y-axis maximum and minimum values are within brackets.

**Figure S3.**
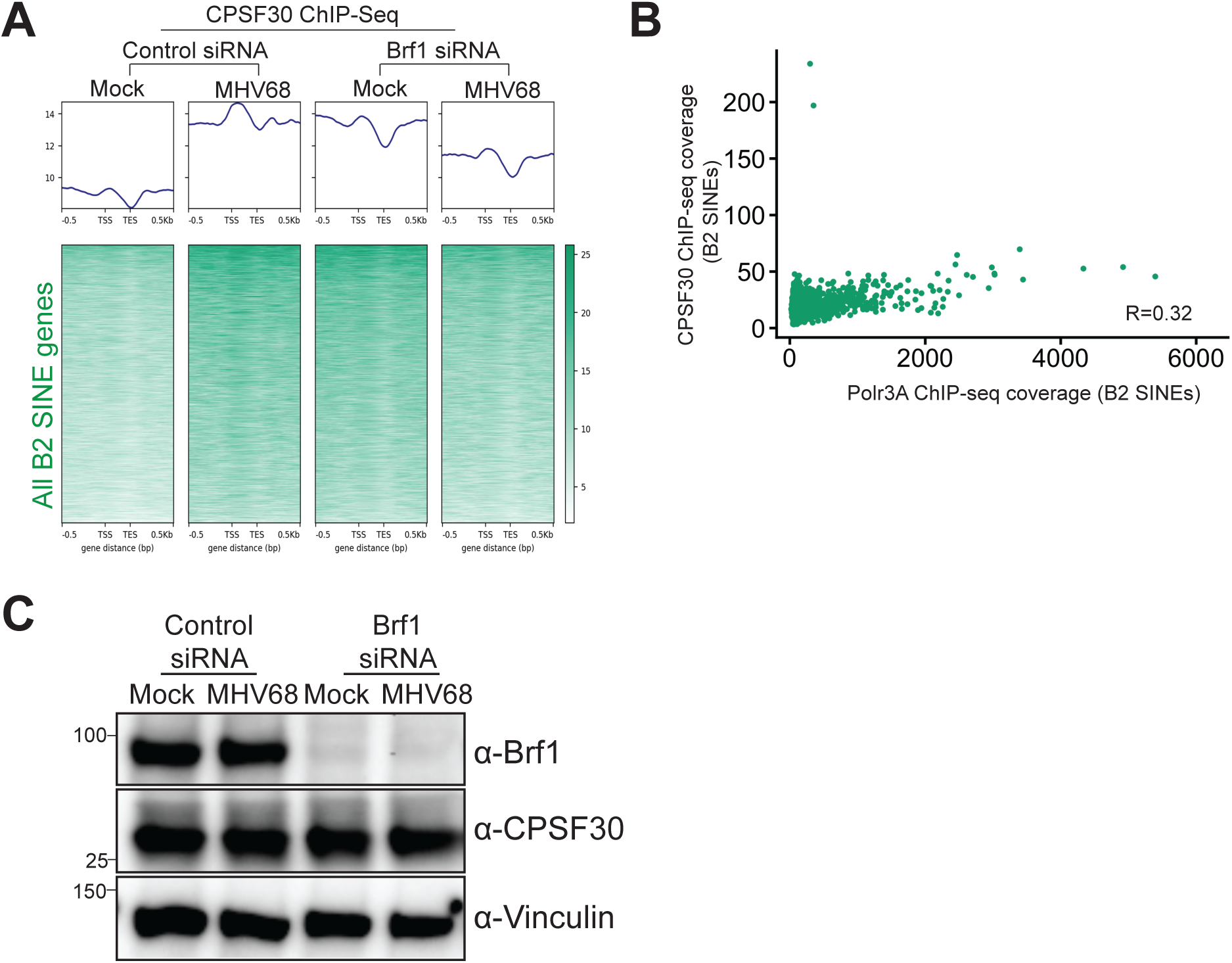
CPSF30-seq analysis at B2 SINE loci during MHV68 infection (A) NIH3T3 cells transfected with control non-targeting or Brf1 targeting siRNAs were mock-treated or infected with MHV68 at an MOI of 5. At 24hpi, cells were harvested for ChIP-seq. The top panels are metagene plots displaying CPSF30 ChIP-seq signal across B2 SINE genes (n=3434) with detectable Polr3A peaks (from Fig.1) and expressed during infection (RPKM≥5) (4). Alignment files were averaged from two replicates. ChIP-seq signal was plotted as a histogram with 10 bp bins for −500 to +500bp around the transcription start site (TSS). In the bottom panel, each row of the heat map corresponds to the CPSF30 ChIP-seq signal for each B2 SINE gene from the metaplot above. (B) Polr3A ChIP-seq signals associated with Polr3A occupied peaks at B2 SINEs during infection (from Fig. 1) were plotted against CPSF30-ChIP-seq signals. The correlation (R) between ChIP-seq signals is denoted on the graphs and was calculated using the Pearson correlation coefficient. (C) NIH3T3 murine fibroblasts transfected with control non-targeting or Brf1 targeting siRNAs were mock-treated or infected with MHV68 at an MOI of 5. At 24hpi, cells were harvested and lysed to extract total protein and were analyzed by Western blotting with antibodies against Brf1, CPSF30, and Vinculin (loading control).

**Figure S4.**
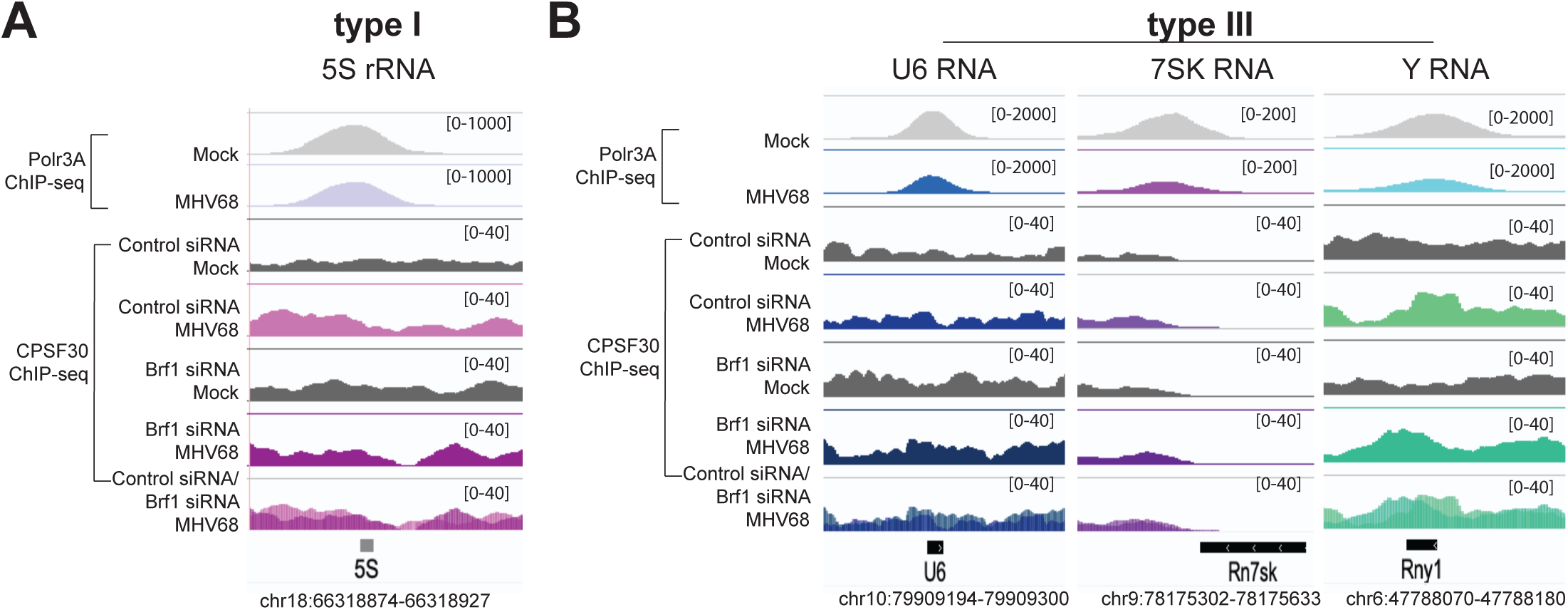
CPSF30 ChIP-seq signal at type I and type III promoters on the cellular genome Polr3A ChIP-seq coverage across select promoter type I (A) and type III (B) genes from mock-treated or MHV68-infected samples. Alignment files were averaged from two replicates and visualized in the Integrative Genome Viewer (3). Y-axis maximum and minimum values are within brackets.

---

Dataset S1 (separate file). Polr3A ChIP-Seq Differential Binding Analysis (MHV68 vs. Mock). DiffBind (5) was used to perform differential binding analysis of Polr3A occupancy data from mock-treated and MHV68-infected samples. The values shown in the dataset are log_2_ fold change (FC) (MHV68 vs. Mock) and the false-discovery rate (FDR) *p*-value at locations within the genome found to have peaks using MACS2 (1).

Dataset S2 (separate file). B2 SINE - ChIRP coupled to mass spectrometry (ChIRP-MS). NIH3T3 cells were mock-treated or infected with MHV68 at an MOI of 5. Comprehensive identification of RNA-binding proteins (ChIRP) was performed using biotinylated B2 SINE oligos. Antisense LacZ oligos were used as a negative control. Protein was extracted from ChIRP samples and was subjected to mass-spectrometry. The dataset shows peptide counts and percent coverage detected in the samples.

## Notes

https://github.com/UCB-GLab/SAMBAR-Net

https://www.ncbi.nlm.nih.gov/geo/query/acc.cgi?acc=GSE288677

### Competing Interest Statement

The authors have declared no competing interest.

